# Neural Tracking of the Maternal Voice in the Infant Brain

**DOI:** 10.1101/2025.04.01.646597

**Authors:** Sarah Jessen, Martin Orf, Jonas Obleser

## Abstract

Infants preferentially process familiar social signals, but the neural mechanisms underlying continuous processing of maternal speech remain unclear. Using EEG-based neural encoding models based on temporal response functions, we investigated how 7-month-old human infants track maternal vs. unfamiliar speech and whether this affects simultaneous face processing. Infants (13 boys, 12 girls) showed stronger neural tracking of their mother’s voice, independent of acoustic properties, suggesting an early neural signature of voice familiarity. Furthermore, central encoding of unfamiliar faces was diminished when infant’s heard their mother’s voice and face-tracking accuracy at central electrodes increased with earlier occipital face tracking, suggesting heightened attentional engagement. However, we found no evidence for differential processing of happy vs. fearful faces, contrasting previous findings on early emotion discrimination. Our results reveal interactive effects of voice familiarity on multimodal processing in infancy: while maternal speech enhances neural tracking, it may also alter how other social cues, such as faces, are processed. The findings suggest that early auditory experiences shape how infants allocate cognitive resources to social stimuli, emphasizing the need to consider cross-modal influences in early development.

**Significance statement:** How infants continuously process familiar social signals and how this familiarity shapes perception has remains unknown. Here we demonstrate that infants’ brains preferentially track maternal speech over unfamiliar voices, highlighting the early tuning of the auditory system to socially relevant signals. Moreover, the maternal voice modulates the neural encoding of concurrently presented faces without eliciting emotion-specific differences. These findings underscore the role of caregiver signals in shaping multisensory integration during early development.

## Introduction

Infants are born into a social environment and, within their first year, develop a range of abilities which allow them to interact with their environment with increasing competence.

One fundamental aspect is the ability to recognize familiar people, most importantly their primary caregivers. A number of studies suggest that newborns quickly learn to recognize their mother’s face (Sai, 2005) and voice (DeCasper & Fifer, 1980), and in the first months of life show a preference for such maternal signals (DeCasper & Fifer, 1980; Sai, 2005). Furthermore, social signals like speech elicit differential brain activation when originating from the mother versus an unfamiliar person (Dehaene-Lambertz et al., 2010; Naoi et al., 2012).

While these studies suggest a preferential orientation towards the maternal voice, it is unclear (a) whether infants preferentially process maternal speech over a longer duration and (b) whether this affects other social processes. To investigate these questions, we examined neural tracking of a continuous stream of maternal vs. unfamiliar speech. Neural tracking reflects the neural representation of a continuous signal, and is increasingly used to investigate infant speech processing (Jessen et al., 2019; Kalashnikova et al., 2018; Menn et al., 2022; Ortiz Barajas et al., 2021).

In particular, several studies have investigated which features of an auditory signal infants track most effectively. For example, infants show stronger tracking of infant-directed versus adult-directed speech (Kalashnikova et al., 2018; Menn et al., 2022) and intelligible versus unintelligible speech (Florea et al., 2024). Furthermore, infants’ neural tracking reflects their attunement to their native language; while newborns show equal tracking of familiar and unfamiliar language input, by 6 months, they show weaker amplitude but not phase tracking of their native language (Ortiz Barajas et al., 2021). Auditory tracking in infants is also found for variants of speech input, such as songs (Nguyen et al., 2023) and nursery rhymes (Attaheri et al., 2022), where increased tracking can be found depending on song type (Nguyen et al., 2023). In summary, most previous studies compare infants’ tracking of speech signals with different acoustic properties (infant-directed speech vs. adult-directed speech, familiar vs. unfamiliar language, play song vs. lullaby).

However, maternal and unfamiliar speech do not systematically differ in acoustic properties; the same speech signal can be familiar to one infant and unfamiliar to the next. Hence, potential differences in neural tracking cannot arise from low-level stimulus properties but must arise from familiarity with the voice as such, or from increased attention due to personal significance. While in adults, voice familiarity indeed enhances neural tracking (Yahav et al., 2024), it is unknown whether similar effects can be observed in infancy.

In addition to investigating the differential processing of maternal compared to an unfamiliar voice, we were interested in whether hearing the mother’s voice influences unrelated social processes, as seen with other signals of maternal presence. Recent work for instance shows that maternal odor influences face categorization (Leleu et al., 2020) and emotional face processing (Jessen, 2020). However, social odor is a more subtle social signal, and often not consciously processed (Loos et al., 2025). Hence, hearing the mother’s voice might be more attention grabbing, potentially interfering with the processing of other social signals. For instance, it has been observed in adults that hearing a task-irrelevant voice in the background impairs face memory (Bell et al., 2019).

To investigate these questions, we studied 7-month-old infants in two sessions. In one, they listened to a recording of their mother reading a children’s story; in the other, they heard another infant’s mother reading the same story. While listening, infants viewed photos of happy and fearful facial expressions. We analyzed event-related potentials (ERPs) linked to attention (Nc; Guy et al., 2016; Xie et al., 2019) and face processing (N290, P400; Guy et al., 2016; Xie et al., 2019), as well as neural tracking of faces and voices, which allowed us to directly link voice and face tracking.

## Methods

### Pre-registration

Rationale, sampling plan, and major analyses were pre-registered prior to data acquisition at https://aspredicted.org/x4xc-rwbt.pdf. In what follows we will refer to this and indicate which analyses deviated from the pre-registered analyses.

### Participants

A total of 39 7-month-old infants were invited to participate in the study. Five additional infants participated only in the first appointment and were therefore excluded from further analysis. The exact composition of the analysis sample differs for the two analysis tracks we will present (see below).

Infants were recruited via the maternity unit of the local hospital (Universitätsklinikum Schleswig-Holstein). All infants were born full-term (at least 37 weeks gestational age) with a birthweight of at least 2500 g, and had no known neurological deficits or visual or hearing impairments. The study was approved by the local ethics committee, conducted according to the Declaration of Helsinki, and parents provided written informed consent prior to the infant’s participation.

For the evoked-potentials averaging (ERP) analysis, the final sample included N = 25 infants (age at first appointment: 214 ± 7 days (mean ± standard deviation), age at second appointment: 223 ± 8 days; 13 boys, 12 girls). The remaining 14 infants were excluded from the analysis because they did not contribute at least 10 trials per Emotion condition at one of the appointments (n= 9); had an overall mean Nc amplitude more than 2 standard deviations from the mean (n=3); or continued to cry at one of the appointments (n=2).

For the encoding-model analysis using multivariate temporal response functions (mTRF), the final sample included N = 30 7-month-old infants (15 boys, 15 girls, age at first appointment: 214 ± 7 days, age at second appointment: 221 ± 7 days). The remaining 9 infants were excluded because they did not contribute at least 100 artifact-free data segments at one of the appointments (n=6); continued to cry at one of the appointments (n=2); or because the trigger for the sound onset was not recorded (n=1). All but one infant included in the ERP analysis were also included in the mTRF analysis.

We aimed for a sample size of N = 25 in the ERP analysis based on our pre-registration. A sample size of N = 25 allowed us to detect effects of at least medium size (f =.25) in a 2 × 2 within-subjects ANOVA with a typical type-I error rate of.05 with a power of.8 (assuming a correlation of.5 of the repeated measures, G*Power 3.1), which seemed sufficient given the large effect observed in Jessen (2020) for the interaction between *Emotion* and *Familiarity* in a very similar paradigm (f =.39).

### Stimuli

As visual stimuli, we used photographs of Caucasian female faces displaying happy and fearful facial expressions from the FACES database ((Ebner et al., 2010), ID 54, 63, 85, 90, 115, 173). Faces were cropped to an oval shape so that as little of the hair as possible was visible.

As auditory stimuli, we used an approximately one-minute-long recording of a children’s story, read and recorded by each infant’s mother. The story was identical for all participants and the recording was made at the first appointment prior to the actual experiment. The mother was given a printout of the story and instructed to read the story “as if you were reading it to your child”. No other restrictions or instructions were given regarding speaking style or speed. The recording was made via a standard headset using Audacity and normalized to –23 RMS dBFS (dB full scale) after recording. Each audio recording was used twice in the study: Once as a familiar stimulus for the mother’s own infant and once as an unfamiliar stimulus for a different infant.

### Experimental Design

To investigate the impact of a familiar voice on the processing of emotional faces, we followed a 2 × 2 design with the factors *Emotion* (happy, fearful) and *Voice* (familiar, unfamiliar), in which infants heard a voice over loudspeakers while simultaneously viewing photographs of faces presented on a screen (Figure 1).

**Figure 1.**
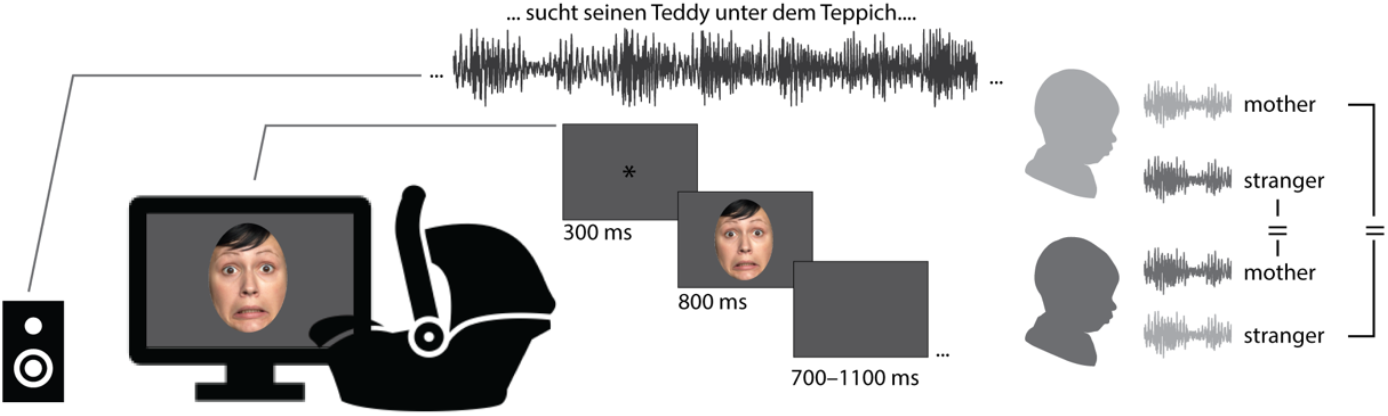
Experimental Set-up and Design. Infants were seated in a car seat in front of a screen with speakers on both sides of the screen. While the recording of the mother or the stranger reading the same story was played continuously in the background, infants viewed photographs of happy and fearful faces. Each recording was used twice in the study; once as a familiar voice for the mother’s own child and once as a stranger’s voice for a different child.

Infants were invited to two appointments, ideally within two weeks. At one appointment, infants would hear their mother’s voice, while at the other appointment, they would hear the unfamiliar voice of a different infant’s mother. The order of *Familiarity* condition was counterbalanced across infants.

Otherwise, the experimental design was identical for both appointments. A maximum of 216 pictures were presented (18 per emotion and actress) arranged in blocks of 24 pictures (12 happy, 12 fearful), which were played consecutively without interruption. Pictures were presented in a pseudorandomized order, ensuring that no picture was repeated more than once. Each picture was presented for 800 ms, preceded by a fixation cross shown for 300 ms and followed by an intertrial interval jittered between 700 and 1100 ms. The audio recording (familiar or unfamiliar voice, depending on the appointment) was played without interruption in a loop throughout the entire experiment in parallel to the visual presentation.

### Procedure

Prior to the first appointment, parents were sent a set of questionnaires which they were asked to complete at home and bring along to the lab appointment (EPDS (Cox et al., 1987), IBQ-R (Gartstein & Rothbart, 2003; Vonderlin et al., 2012), and a lab-internal questionnaire assessing demographic information as well as feeding and sleeping routines of the infant). Questionnaire data are not considered further in the present analysis.

Upon arrival at the lab, parents and infants were familiarized with the new environment and parents were again explained the procedure and had the opportunity to ask any remaining questions, before signing the informed consent form. The EEG recording was prepared while the infant was sitting on their parent’s lap. An elastic cap (BrainCap, Easycap GmbH) containing 27 AgAgCl electrodes according to the international 10-20 system was used for recording. The EEG signal was recorded at a sampling rate of 500 Hz using a BrainAmp amplifier and the software BrainVisionRecorder (both BrainProducts).

During the experiment, the infant was seated in an age-appropriate car seat (Maxi Cosi) placed on the floor approximately 70 cm in front of a 24-inch monitor on which the visual stimuli were presented. The face height was approximately 28 cm. Auditory stimuli were presented via loudspeakers positioned left and right of the monitor set at the same volume for all participants. Matlab (The MathWorks, Inc., Natick, MA) and the toolbox Psychtoolbox (Brainard, 1997) were used to run the experiment. The parents remained in the room with the infant but were instructed to maintain a distance of at least 1 m behind the infant.

The infant was observed and videotaped throughout the experiment by a small camera mounted on top of the monitor. If necessary, the experimenter could interrupt the experiment to play short video clips containing colorful moving shapes and ringtones to redirect the infant’s attention to the screen. The experiment continued until all 216 trials had been presented (approximately 8 minutes) or the infant became too fussy to continue the experiment.

### Data analysis

EEG data were analyzed using Matlab 2023b (The MathWorks, Inc., Natick, MA), using custom-made scripts was well as the toolbox FieldTrip (Oostenveld et al., 2011). Jamovi (version 2.6.26) was used for the correlation analysis and mixed models for the ERP results using the GAMLj package (version 2.4.0).

As preregistered, we analyzed the data focusing on differences in the ERP response to emotional faces, neural tracking of familiar vs. unfamiliar voices, as well as the relation between both measures. Exploratorily and in addition to the preregistered analyses, we also included the responses to emotional faces in the encoding model.

### EEG preprocessing

In a first step, we computed an independent component analysis (ICA) to identify artifactual components. To do so, data were offline re-referenced to the average reference; filtered using a 1-Hz highpass and 40-Hz lowpass filter; and segmented in 1-second epochs. Epochs in which the standard deviation exceeded 100 μV in a sliding window of 200 ms at any electrode were removed from further analysis. On the remaining data, we computed an ICA; on average, 5 ± 2 components (range: 0-9) were identified based on visual inspection of time courses and topographies of the components.

For the ERP analysis, the raw EEG data were filtered between 0.2 and 20 Hz, re-referenced to the linked mastoids; previously identified ICA components were removed; and data were epoched into trials ranging from 200 ms before to 800 ms after the picture onset. Any trials in which the standard deviation exceeded 80 μV in a sliding window of 200 ms at any electrode were removed, and the data were inspected visually for any remaining artifacts. Finally, trials in which the infants did not attend to the screen based on the video recording were removed from further analysis.

Infants who contributed fewer than 10 trials per condition were excluded from further analysis (n=9). Infants in the final sample contributed an average of 37 ± 19 trials per condition (mother-happy: 34 ± 18; mother-fearful: 33 ± 16; stranger-happy: 42 ± 22; stranger-fearful: 41 ± 20; since infants contributed more trials when they heard a stranger’s as opposed to their mother’s voice [F(1,24) = 3.99, p=.057], we exploratorily included number of trials as a covariate in the GLM, which had no effect on the observed results).

For the neural encoding-model analysis, artifactual ICA components were removed and data were filtered between 1 and 10 Hz, but no further visual inspection was carried out (Fiedler et al., 2019; Jessen et al., 2019; Orf et al., 2023).

### Encoding model analysis

A temporal response function (TRF) serves as a simplified model of brain activity, illustrating how the brain processes a given stimulus feature to generate the observed EEG signal, assuming a linear filter mechanism. To compute the TRF, we utilized a multiple linear regression approach (Crosse et al., 2016). To predict the recorded EEG response, we trained a forward model per EEG channel using the envelopes of the mothers and stranger streams (e.g., Fiedler et al., 2019). In this framework, we examined delays between envelope changes and brain responses of between -200 and 800 ms.

To account for EEG variance associated with the processing of presented faces and their corresponding evoked responses, we incorporated triggers for face presentations as regressors using stick functions. A stick function is a modeling vector in which non-event times are denoted by zeros, while events, such as the onset of a face, are marked with a value of one.

To prevent overfitting, we used ridge regression to estimate the TRF and determined the optimal ridge parameter through leave-one-out cross-validation for each participant. We predefined a range of ridge values, calculated a separate model for each value, and averaged across trials to predict the neural response for each test set. The ridge parameter with the lowest mean squared error (MSE) was selected as the optimal value specific to each subject. One model was trained using predictor variables for the envelopes of mother and stranger streams, as well as stick functions for presented faces (Orf et al., 2023). Hence, the model contained three regressors: the speech envelope (which was either the mother’s or the stranger’s voice), one regressor coding for the onset of the happy faces, and one regressor coding for the onset of the fearful faces. These regressors were modelled jointly (same regressor matrix) using the same regularization.

Neural speech tracking quantifies the representation of a single stream within the EEG signal by using TRFs to predict the EEG response. The neural tracking metric (r) was computed by correlating the predicted EEG response with the actual EEG signal using Pearson correlation. A leave-one-out cross-validation approach was used to predict the EEG signal. To assess neural tracking accuracy over TRF time lags, we used a sliding time window (48 ms, 6 samples, 24 ms overlap), resulting in a time-resolved measure of neural tracking (Fiedler et al., 2019; Hausfeld et al., 2018; Kraus et al., 2021; O’Sullivan et al., 2015).

### ERP analysis

For the ERP analysis, we computed single-subject averages for all four conditions (mother-happy, mother-fearful, stranger-happy, stranger-fearful).

We computed the Nc response as the mean value in a time-window of 400–800 ms after picture onset averaged across F3, Fz, F4, C3, Cz, C4, and the N290 and P400 as the mean value in a time-window of 200–300 ms respectively 300–500 ms after picture onset averaged across O1 and O2. These time-windows and electrodes correspond to those preregistered and are based on prior studies (Jessen, 2020; Quadrelli et al., 2019). Exploratorily, we also computed the Nc in a time-window from 350 to 600 ms, as inspection of the average across all conditions revealed an Nc peak in this earlier time-window.

Participants whose Nc amplitude was more than 2 standard deviations from the average across all conditions between 400 and 800 ms were excluded from further analysis (n=3).

### Statistical Analysis

We statistically evaluated both, conventional ERP results and encoding model outcomes, using linear mixed-effects models. These models account for the hierarchical structure in multi-condition, within-subject designs by combining fixed effects and random effects. Predictors were centered and scaled as needed. Random intercepts were included where appropriate to account for inter-individual differences.

The mTRFs were analyzed in Matlab using the fitlme function and the following model:

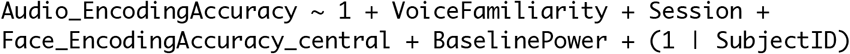

where *VoiceFamiliarity* was the voice heard (mother or stranger). We additionally included *Session* (1 or 2) to account for potential order effects and *SubjectID* to account for interindividual variance. Furthermore, *BaselinePower* (log-transformed) served as a measure of noise in the infant data, while *Face_EncodingAccuracy_central* at central electrodes (C3, Cz, C4) was included to control for the impact of the visual stimuli (faces viewed by the infant) on encoding accuracy.

We also computed the following model to the assess impact of voice encoding on central face processing:

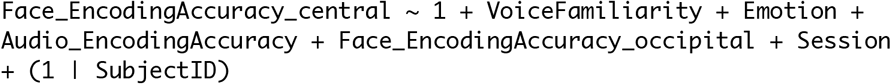

Here, we additionally included *Emotion*, accounting for the emotion expressed in the picture seen (happiness or fear) and face encoding at occipital electrodes (O1, O2) in an earlier, common time window (200 – 300 ms) as *Face_EncodingAccuracy_occipital* to account for visual processing. Including BaselinePower in the model did not further improve the model fit. Not shown in the pseudo-formula above are all two-way interaction terms of the regressors of interest, which we included in the model as well.

We analyzed the ERP data in Jamovi using the following model for each ERP component separately:

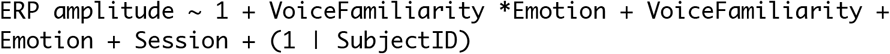

Models additionally including *Trialnumber* (reflecting the number of trials in each condition) and/or *Breastfeeding* (whether the infant was still breastfed or not) yielded the same results as reported below.

Finally, to link our two EEG analysis approaches and for completeness, we also report a simple Pearson correlation between *Audio_EncodingAccuracy* and the ensuing difference in Nc response to happy and fearful faces.

For the pre-registered hypotheses (see below) and substantive null effects of interest (i.e., the absence of effects of Emotion in face encoding models), we report Bayes Factors in favor of the null effect derived from the Jamovi model output using Eq. 6 in Faulkenberry (2018) (see also Wagenmakers, 2007).

### Data availability

Upon publication, data and anaylsis scripts will be made available at http://osf.

## Results

### Infants’ neural speech tracking is sensitive to voice familiarity

The temporal response function showed clear signatures of the infant brain neurally tracking both, the mother’s as well as the stranger’s voice. The frontocentral bilateral distribution (Figure 3A) of this neural tracking is expected and well in line with the adult neural tracking literature, broadly commensurate with bilateral auditory (i.e., perislyvian cortical) origins.

However, tracking of the mother’s voice was consistently stronger than tracking of a stranger’s voice (Figure 3), despite both having comparable acoustic properties (Figure 2). This is highlighted in Figure 3C, which compares the encoding accuracy within the time window with the highest TRF and time-shifted neural tracking (200–300 ms). A statistically significant enhancement in neural tracking for the mother’s voice compared to the stranger’s voice is evident (b = -0.004, SE = 0.001, t(84.8) = -4.72, p <.001).

**Figure 2.**
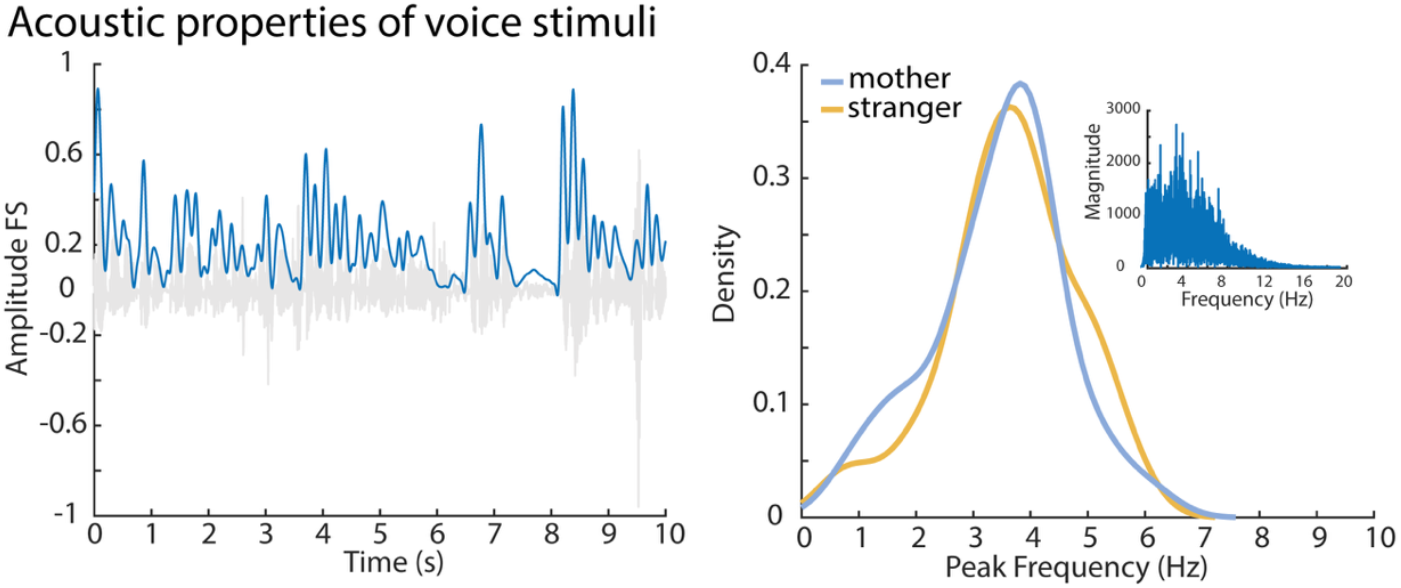
Acoustic properties of voice signals. As regressor for auditory neural tracking, we used the voice envelope, as shown in the left panel. The envelope was extracted from both the mother and stranger condition audio signal, and the peak frequency was identified from the spectrum using the Fast Fourier Transform (FFT). Since the same recordings were used once as a stranger and once as a familiar (mother’s) voice, they showed very similar acoustic properties, as can be seen in the right panel. Histograms and Kernel Density Estimates (KDE) with a bandwidth of 0.5 were used to visualize the distribution of peak frequencies across subjects for both conditions. (Note that the two stimulus distributions are not identical as not all infants could be included in the final sample, so that for a small number of infants the recording was only included in one of the conditions).

**Figure 3.**
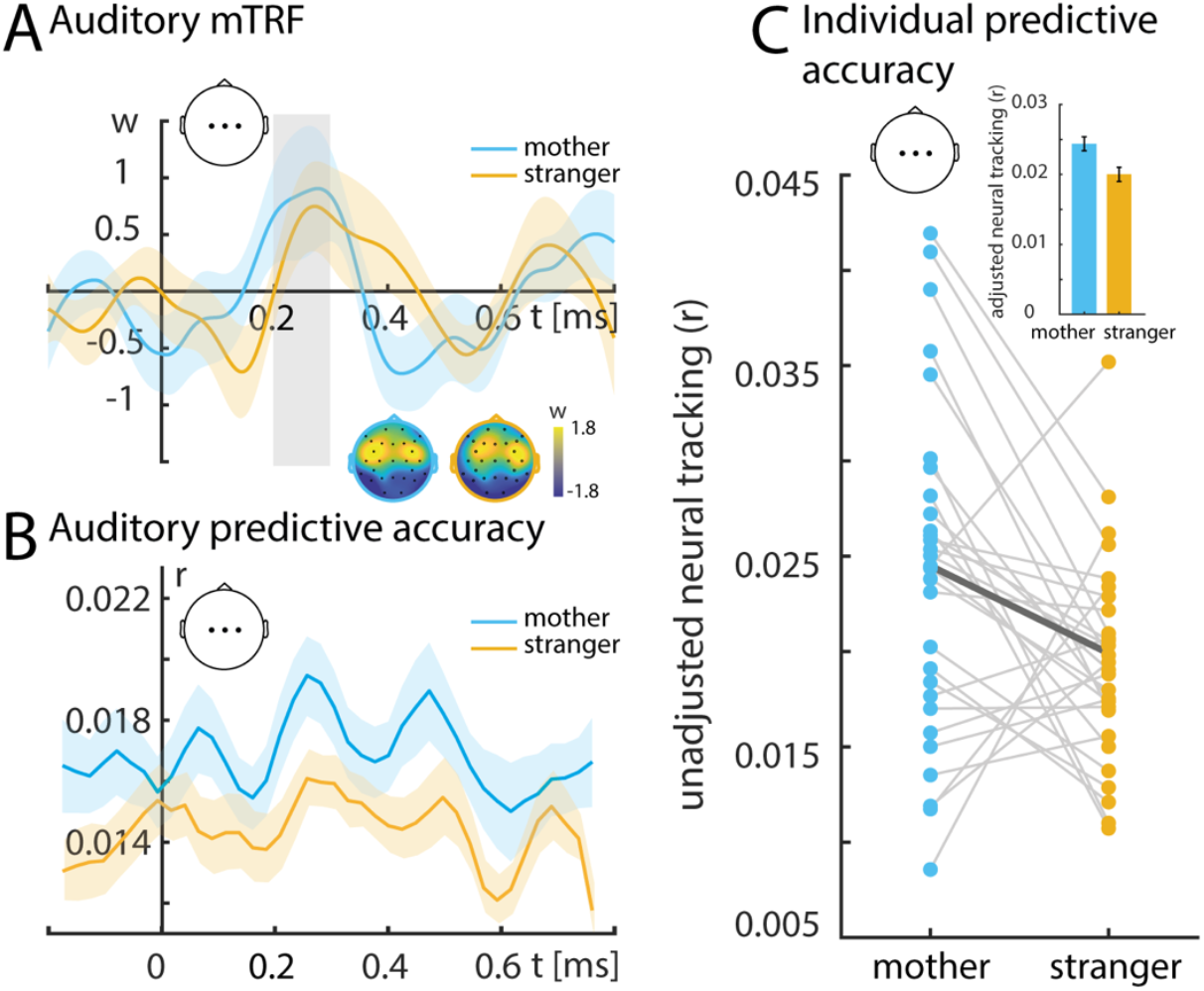
Temporal response functions (TRFs) and neural speech tracking. **A**. TRF weights are averaged across infants and channels of interest (Cz, C3, C4). The shaded areas show the standard error for each time lag across infants. The inset displays the topographical distribution of TRF weights for a time window between 200-300 ms (gray bar). Blue colors indicate TRFs for the mother’s voice, while orange colors indicate TRFs for the stranger’s voice. **B**. Neural tracking unfolds over time lags (-200 to 800 ms). Solid lines show the averaged predictive accuracy (r) across subjects and channels of interest (topographic map). Shaded areas represent the standard error for each time lag across infants. **C**. Unadjusted neural tracking (r) refers to the encoding accuracy (200-300 ms), calculated by multiplying the estimated TRFs by the presented audio envelope. The line plot shows single-infants data (N = 30), averaged over channels of interest (Cz, C3, C4). Connecting lines between dots indicate the same subject. Blue dots represent tracking of the mother’s voice, while orange dots represent tracking of the stranger’s voice. The inset shows adjusted neural tracking based on predictive model results, including covariates such as baseline power and face tracking.

### Infant encoding models of faces are sensitive to voice familiarity

Furthermore, we clearly observed neural tracking of faces at both, central and occipital electrodes (i.e., face encoding accuracies greater than zero; Figure 4). This neural face-encoding, however, was modulated by the concurrent presence or absence of the mother’s familiar voice.

**Figure 4.**
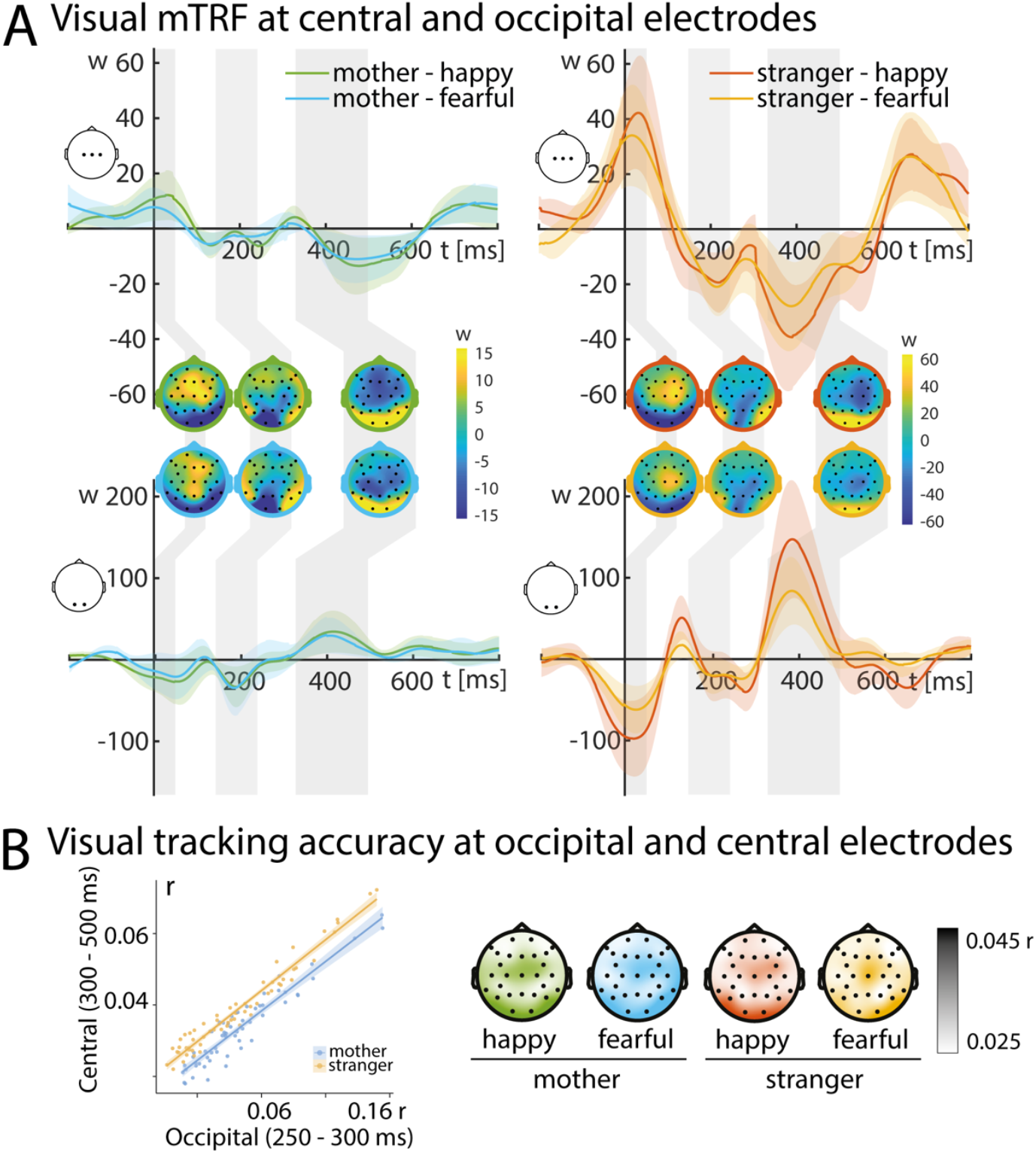
Temporal response function and neural face tracking. **A**. Clear tracking was found at central (C3, Cz, C4) and occipital (O1, O2) electrodes, which was lower when infants heard their mother (left panel; blue/green) compared to a stranger (right panel; red/ orange). We did not find an impact of face emotion, suggesting that happy (green/red) and fearful faces (blue/orange) were tracked to a similar extent. Faces were encoded at both, central and occipital electrodes, as can be seen in the topographic representations (350 - 500 ms). **B**. Encoding at central electrodes (300 – 500 ms) was modulated by both, voice identity (blue = mother, orange = stranger) and preceding occipital encoding (250 – 300 ms). Central face encoding was high when a stranger’s voice was heard (p =.004, see text for details), and was independently modulated by preceding occipital face encoding, p < 0.001; interaction n.s.). As can be seen from the topographic representations, differences where focused on central and occipital electrodes as can be expected for face processing. Topographical representation show face encoding accuracy between 350 – 500 ms.

When infants heard a stranger as opposed to their mother, central electrodes showed an enhanced face encoding (Effect of *VoiceFamiliarity* on *Face_EncodingAccuracy_central*; b= 0.008, SE = 0.003, t(90.9) = 2.98, p=.004). This can also be expressed as a relatively diminished encoding of (unknown) faces when the mother’s voice is present (Figure 4A).

Independent of the heard voice identity, the strength of an infant’s central face tracking was robustly predictable from a preceding encoding accuracy at occipital electrodes (*Face_EncodingAccuracy_occipital*; b=0.010, SE = 0.001, t(105.9) = 6.67, p<.001). This lends plausibility to the overall results of the forward encoding model (which establishes encoding accuracies per electrode): It confirms that infants who showed stronger tracking at occipital electrodes in an earlier time window also showed stronger tracking at central electrodes.

### Absence of facial emotion effects

We did not find an effect of *Emotion* expressed in the face stimuli on the temporal response function or the resulting encoding accuracy at either central or occipital electrodes (both Bayes Factors >10 in favour of the null hypothesis). Neither of the *Emotion* differences discernable in some time windows and electrodes did surpass conventional levels of statistical significance. This speaks to a generally reduced impact of *Emotion* on infants’ neural responses in this paradigm with a spoken-language backing track, compared to many previous infant studies on face processing without such auditory input.

### Infant evoked responses to faces do not yield effects of voice familiarity or emotion content

We did not find significant main effects of *VoiceFamiliarity* or *Emotion*, nor an interaction between the two on any of the ERP components investigated (all p>. 1, all BF10 <.35; see Figure 5 for ERP response at central and occipital).

**Figure 5.**
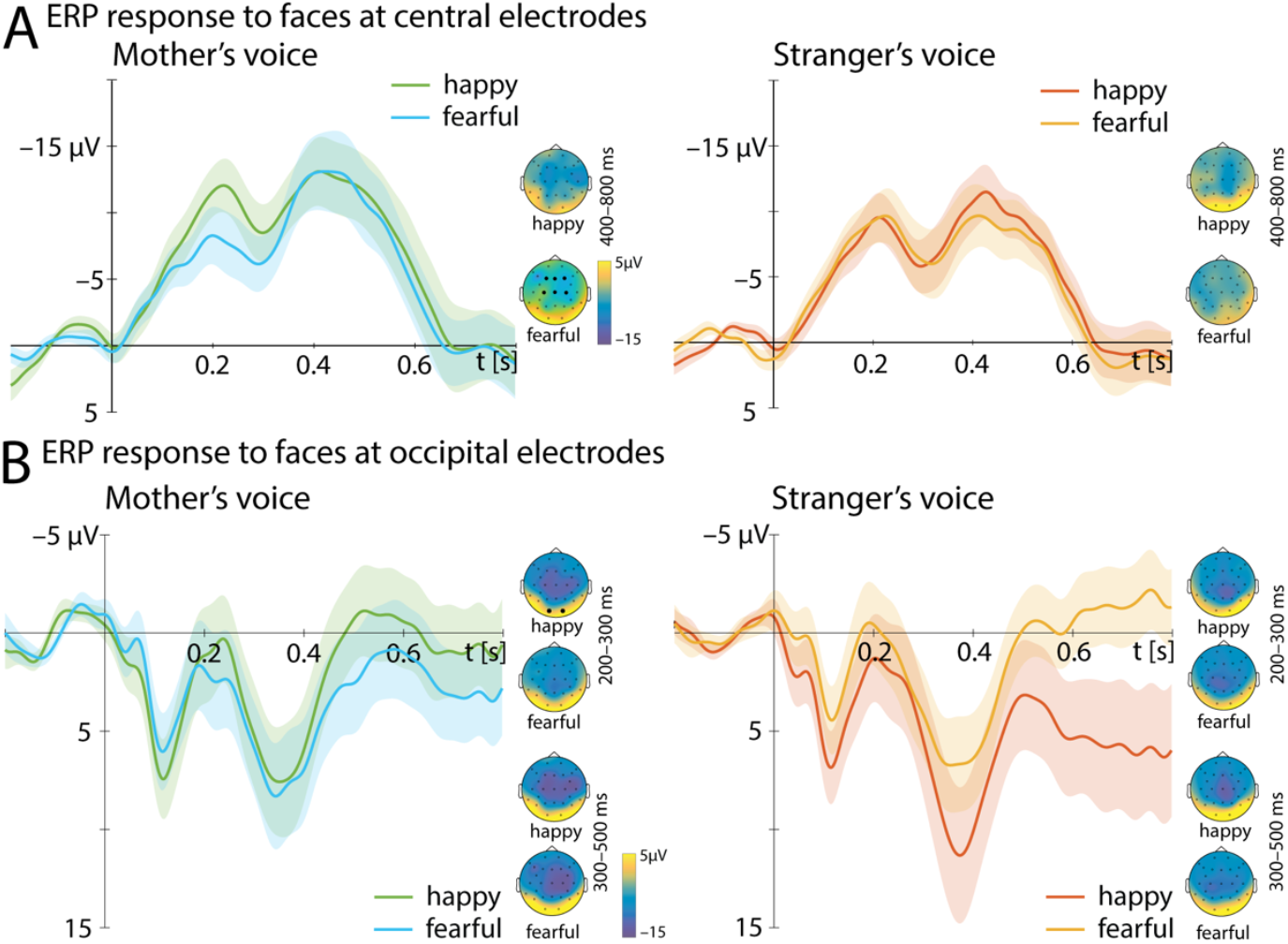
Conventional evoked-response analysis to faces. Shown are ERP responses at central (A) and occipital electrodes (B) to the presentation of happy (lighter colors) and fearful faces (darker colors) while infants listened to either their mother’s voice (left panel, blue/green) or a stranger’s voice (right panel, orange/red). Topographic representations show the average for time-windows analyzed for ERP responses (A: Nc response in a time-window of 400 to 800 ms; B: N290 in a time-window of 200 to 300 ms and P400 in a time-window of 300 to 500 ms. Dots indicate the electrodes averaged (A: F3, Fz, F4, C3, Cz, C4; B: O1, O2). No significant differences were observed.

The difference in Nc response to fearful and happy faces, as obtained in the conventional ERP analysis and often reported in infant ERP studies, was mildly yet non-significantly positively correlated with the strength of auditory tracking of the ongoing voice, as derived from the encoding model (r=.24, p=.10).

## Discussion

Can we trace infants’ neural processing of familiar versus unfamiliar voices and its potential impact on the processing emotional faces? In doing so, we have leveraged neural encoding models of the infant EEG and have contrasted them with conventional evoked-response analyses (ERP).

Infants showed a stronger neural tracking of their mother’s compared to a stranger’s voice. Furthermore, infants’ face processing was affected by the concurrently heard voice; infants hearing a stranger showed a stronger increase in central face tracking. In addition, central face tracking increase with previous occipital tracking. While infants thus successfully encode faces, we found no difference in processing between fearful and happy faces.

### Neural encoding of maternal versus unfamiliar voices

The robust tracking of voice information and the enhanced tracking of maternal over an unfamiliar voices adds to a growing body of literature investigating the continuous tuning of the developing brain to relevant auditory social signals (Jessen et al., 2019; Kalashnikova et al., 2018; Menn et al., 2022; Nguyen et al., 2023).

For the first time, we show that enhanced tracking occurs not only for auditory signals that are either specifically tailored to infants (e.g., infant-directed speech or nursery rhymes, (Kalashnikova et al., 2018; Menn et al., 2022; Nguyen et al., 2023)) or native language input (Ortiz Barajas et al., 2021), but also for voices carrying distinct signals for an individual infant. This enables the infant to attend to personally relevant social cues, facilitating bonding and social learning.

One explanation for stronger tracking of maternal voice is better predictability due to the high familiarity. Such an explanation would be comparable to the mechanisms observed in the case of infant-directed speech or lullabies, both of which are characterized by a high degree of rhythmicity, large modulations in pitch, and simplified structure, all features that increase predictability and thereby facilitate processing (Kalashnikova et al., 2018). For maternal speech, increased predictability is more subtle and individualized, resulting from the idiosyncrasies of each speaker. A similar effect is seen in adults, where speaker familiarity improves comprehension for hearing-impaired listeners (Souza et al., 2013) and even brief familiarization benefits listeners with intact hearing (Nygaard & Pisoni, 1998).

While such an explanation is primarily based on perceptual facilitation, another explanation could be derived from attention-related processes. Since maternal signals are highly salient and of particular importance to the infant, infants allocate more attentional resources to maternal signals (DeCasper & Fifer, 1980; Rigato et al., 2023). As has been extensively studied in adults, increased attention allocation leads to increased neural tracking (e.g., Fiedler et al., 2019; Mesgarani & Chang, 2012; Orf et al., 2023; Reetzke et al., 2021), which is likely to be the case also in infancy.

### Neural encoding of faces is influenced by voice familiarity

While an increasing number of studies investigate auditory tracking, visual tracking is less commonly investigated. However, initial studies suggest neural tracking also of continuous visual signals (Jessen et al., 2019; Ki et al., 2021). In our study, we did not investigate the tracking of a continuous visual signal but of discrete visual events optimized for an ERP design. Nevertheless, we observed clear visual tracking at both central and occipital electrodes, corresponding in both, topography and latency, to well-known infant ERP components typically observed in face processing (occipital N290-P400 and central Nc; Conte et al., 2020; Xie et al., 2019). This observation is confirmed by the more traditional ERP analysis. Thus, infants successfully encode faces even when listening to unrelated voice information in parallel, and such a processing is observed not only in the ERP response but also in neural tracking.

Moreover, the encoding of faces is not independent of voice processing. Voice information modulated the relation between occipital and central encoding; when hearing an unfamiliar voice, faces were encoded stronger as reflected by central encoding accuracy (Fig. 4). Here, using mTRFs to analyze face processing offers a distinct advantage: Note that these face encoding accuracies result from a forward encoding model that allowed us to directly control for simultaneous auditory tracking within in the same model (e.g., Orf et al., 2023).

While purely speculative at this point, these data might suggest that an unfamiliar voice causes infants to attend more closely to other types of social information. Conversely, maternal presence might reduce the infants’ need to attend to unfamiliar faces (see Jessen, 2020, for a qualitatively similar effect of maternal odor). It is also possible that infants attempt to match the unfamiliar voice to one of the unfamiliar faces, while the familiar mother’s voice clearly does not match any of the faces (since the infants were not presented with pictures of their mother). In such a scenario, an increased face response when hearing an unfamiliar voice might reflect an increased effort to match the voice to one of the faces.

Lastly, and independent of maternal voice, better central encoding of faces in the typical Nc-like time window was preceded by better occipital encoding in the N290 time window (i.e., a “gating effect”; Fig. 4B). While linear models per se can rarely be interpreted causally, these results may provide putative evidence that Nc-like central face tracking processes is ‘gated by’ (i.e., statistically dependent on) the relative strength of a more ‘upstream’ occipital encoding of these faces. The result clearly calls for a more elaborate neuroimaging approach to sensory–occipital vs attentional– frontocentral processes in the infant brain.

In sum, the analyses of neural tracking revealed that infants track faces effectively even while hearing an unrelated voice, and voice familiarity influences how faces are processed

### Absence of emotion effect in facial processing

Contrary to our prediction, we found no evidence that maternal voice influenced emotion processing, nor did we find evidence for emotion processing per se. This is surprising, given the vast body of literature showing that 7-month-old infants reliably discriminate emotions at the neural level (e.g. Aran et al., 2023; Bowman et al., 2022; Leppanen et al., 2007; Peltola et al., 2009; Xie et al., 2019), and our own previous work, which found robust evidence for emotion discrimination in a very similar set-up using the same visual stimuli (Jessen, 2020). At this point, we can only speculate about possible explanations.

Note that the absence of an emotion effect for infants exposed to maternal signals aligns with our previous findings, where infants showed evidence of emotion discrimination only when exposed to social signals from a stranger (Jessen, 2020). Though not significant, on a descriptive level, this is also observed in the present study at later occipital responses, which show emotion-related differences when the infant is hearing a stranger’s voice but not when they are hearing their mother (in both ERP and mTRF).

However, the question remains as to why no significant difference was observed. While infants at this age typically show neural emotion differentiation, not all studies report emotion differentiation in all three ERP components. For example, both Aran et al. (2023) and Xie et al. (2019) only found a differential processing of happy and fearful faces on the N290 but not on the P400 or Nc, while Leppanen et al. (2007) report no difference on the N290 but rather the P400 and Nc. Hence, while emotion processing in infancy has been extensively studied, it is still unclear under which conditions which neural processes are affected by affective content.

One possibility is that while background voices did not disrupt face processing, they may have hindered detailed extraction of emotional cues. To our knowledge, no EEG study so far has investigated emotional face processing with concurrent, unrelated voice signals. Infants might for instance have been distracted by attending to the voice; an indicator of such an effect might be the fact that infants contributed more trials when listening to a stranger versus their mother (although this effect was only marginally significant). Other potential indicators, such as an interaction between auditory encoding accuracy and emotion or a correlation between auditory encoding accuracy and difference in Nc amplitude were not observed. On the contrary, on a descriptive level, we found a non-significant association between increasing auditory encoding accuracy and increasing Nc amplitude for fearful compared to happy faces (r =.24, p =.1), which would suggest an improved discrimination with increasing encoding accuracy.

Future studies are clearly necessary to further explore this issue and examine the robustness of infants’ emotion processing under conditions of potential distraction.

### Conclusion

Infants are exposed to a wide variety of social signals in their daily lives. We investigated the interplay between two different types of signals, voice and faces, providing unrelated information. Infants showed a stronger tracking of their mother’s than of a stranger’s voice and this tracking impacted how well unrelated faces were encoded. Surprisingly, we found no evidence for more sophisticated face processing, such as emotion discrimination. Our results further highlight the potential of using neural tracking to uncover the complex relationship between different types of social processes in early development.

## Acknowledgements

The authors are grateful to the families for their participation. We further would like to thank the German Research Foundation (DFG) for funding to SJ (JE781/3-1) and to JO (JO352/2-2). Generative Artificial Intelligence (DeepL by DeepL; chatGTP by openAI) was used for proofreading, for the content of which the authors retain full responsibility.

## References

Aran, O., Garcia, S. E., Hankin, B. L., Hyde, D. C., & Davis, E. P. (2023). Signatures of emotional face processing measured by event-related potentials in 7-month-old infants. Developmental Psychobiology, 65(2), e22361. 10.1002/dev.22361

Attaheri, A., Choisdealbha, A. N., Di Liberto, G. M., Rocha, S., Brusini, P., Mead, N., Olawole-Scott, H., Boutris, P., Gibbon, S., Williams, I., Grey, C., Flanagan, S., & Goswami, U. (2022). Delta- and theta-band cortical tracking and phase-amplitude coupling to sung speech by infants. Neuroimage, 247, 118698. 10.1016/j.neuroimage.2021.118698

Bell, R., Mieth, L., Roer, J. P., & Buchner, A. (2019). Effects of Auditory Distraction on Face Memory. Sci Rep, 9(1), 10185. 10.1038/s41598-019-46641-7

Bowman, L. C., McCormick, S. A., Kane-Grade, F., Xie, W., Bosquet Enlow, M., & Nelson, C. A. (2022). Infants’ neural responses to emotional faces are related to maternal anxiety. J Child Psychol Psychiatry, 63(2), 152–164. 10.1111/jcpp.13429

Brainard, D. H. (1997). The Psychophysics Toolbox. Spat Vis, 10(4), 433–436. https://www.ncbi.nlm.nih.gov/pubmed/9176952

Conte, S., Richards, J. E., Guy, M. W., Xie, W., & Roberts, J. E. (2020). Face-sensitive brain responses in the first year of life. Neuroimage, 211, 116602. 10.1016/j.neuroimage.2020.116602

Cox, J. L., Holden, J. M., & Sagovsky, R. (1987). Detection of postnatal depression. Development of the 10-item Edinburgh Postnatal Depression Scale. Br J Psychiatry, 150, 782–786. 10.1192/bjp.150.6.782

Crosse, M. J., Di Liberto, G. M., Bednar, A., & Lalor, E. C. (2016). The Multivariate Temporal Response Function (mTRF) Toolbox: A MATLAB Toolbox for Relating Neural Signals to Continuous Stimuli. Front Hum Neurosci, 10, 604. 10.3389/fnhum.2016.00604

DeCasper, A. J., & Fifer, W. P. (1980). Of Human Bonding - Newborns Prefer Their Mothers Voices. Science, 208(4448), 1174–1176. 10.1126/science.7375928

Dehaene-Lambertz, G., Montavont, A., Jobert, A., Allirol, L., Dubois, J., Hertz-Pannier, L., & Dehaene, S. (2010). Language or music, mother or Mozart? Structural and environmental influences on infants’ language networks. Brain Lang, 114(2), 53–65. 10.1016/j.bandl.2009.09.003

Ebner, N. C., Riediger, M., & Lindenberger, U. (2010). FACES--a database of facial expressions in young, middle-aged, and older women and men: development and validation. Behav Res Methods, 42(1), 351–362. 10.3758/BRM.42.1.351

Faulkenberry, T. J. (2018). Computing Bayes factors to measure evidence from experiments: An extension of the BIC approximation. Biometrical Letters, 55(1), 31–43. 10.2478/bile-2018-0003

Fiedler, L., Wostmann, M., Herbst, S. K., & Obleser, J. (2019). Late cortical tracking of ignored speech facilitates neural selectivity in acoustically challenging conditions. Neuroimage, 186, 33–42. 10.1016/j.neuroimage.2018.10.057

Florea, C., Reimann, M., Schmidt, F., Preiss, J., Angerer, M., Ameen, M., Heib, D., Roehm, D., & Schabus, M. (2024). Neural speech tracking in newborns: prenatal learning and contributing factors. bioRxiv. 10.1101/2024.03.18.585222

Gartstein, M. A., & Rothbart, M. K. (2003). Studying infant temperament via the Revised Infant Behavior Questionnaire. Infant Behavior & Development, 26(1), 64–86. https://doi.org/PiiS0163-6383(02)00169-8 Doi 10.1016/S0163-6383(02)00169-8

Guy, M. W., Zieber, N., & Richards, J. E. (2016). The Cortical Development of Specialized Face Processing in Infancy. Child Dev, 87(5), 1581–1600. 10.1111/cdev.12543

Hausfeld, L., Riecke, L., Valente, G., & Formisano, E. (2018). Cortical tracking of multiple streams outside the focus of attention in naturalistic auditory scenes. Neuroimage, 181, 617–626. 10.1016/j.neuroimage.2018.07.052

Jessen, S. (2020). Maternal odor reduces the neural response to fearful faces in human infants. Dev Cogn Neurosci, 45, 100858. 10.1016/j.dcn.2020.100858

Jessen, S., Fiedler, L., Munte, T. F., & Obleser, J. (2019). Quantifying the individual auditory and visual brain response in 7-month-old infants watching a brief cartoon movie. Neuroimage, 202, 116060. 10.1016/j.neuroimage.2019.116060

Kalashnikova, M., Peter, V., Di Liberto, G. M., Lalor, E. C., & Burnham, D. (2018). Infant-directed speech facilitates seven-month-old infants’ cortical tracking of speech. Sci Rep, 8(1), 13745. 10.1038/s41598-018-32150-6

Ki, J. J., Dmochowski, J. P., Touryan, J., & Parra, L. C. (2021). Neural responses to natural visual motion are spatially selective across the visual field, with selectivity differing across brain areas and task. Eur J Neurosci, 54(10), 7609–7625. 10.1111/ejn.15503

Kraus, F., Tune, S., Ruhe, A., Obleser, J., & Wostmann, M. (2021). Unilateral Acoustic Degradation Delays Attentional Separation of Competing Speech. Trends Hear, 25, 23312165211013242. 10.1177/23312165211013242

Leleu, A., Rekow, D., Poncet, F., Schaal, B., Durand, K., Rossion, B., & Baudouin, J. Y. (2020). Maternal odor shapes rapid face categorization in the infant brain. Dev Sci, 23(2), e12877. 10.1111/desc.12877

Leppanen, J. M., Moulson, M. C., Vogel-Farley, V. K., & Nelson, C. A. (2007). An ERP study of emotional face processing in the adult and infant brain. Child Dev, 78(1), 232–245. 10.1111/j.1467-8624.2007.00994.x

Loos, H. M., Schaal, B., Pause, B. M., Smeets, M. A. M., Ferdenzi, C., Roberts, S. C., de Groot, J., Lubke, K. T., Croy, I., Freiherr, J., Bensafi, M., Hummel, T., & Havlicek, J. (2025). Past, Present, and Future of Human Chemical Communication Research. Perspect Psychol Sci, 20(1), 20–44. 10.1177/17456916231188147

Menn, K. H., Michel, C., Meyer, L., Hoehl, S., & Mannel, C. (2022). Natural infant-directed speech facilitates neural tracking of prosody. Neuroimage, 251, 118991. 10.1016/j.neuroimage.2022.118991

Mesgarani, N., & Chang, E. F. (2012). Selective cortical representation of attended speaker in multi-talker speech perception. Nature, 485(7397), 233–236. 10.1038/nature11020

Naoi, N., Minagawa-Kawai, Y., Kobayashi, A., Takeuchi, K., Nakamura, K., Yamamoto, J., & Kojima, S. (2012). Cerebral responses to infant-directed speech and the effect of talker familiarity. Neuroimage, 59(2), 1735–1744. 10.1016/j.neuroimage.2011.07.093

Nguyen, T., Reisner, S., Lueger, A., Wass, S. V., Hoehl, S., & Markova, G. (2023). Sing to me, baby: Infants show neural tracking and rhythmic movements to live and dynamic maternal singing. Dev Cogn Neurosci, 64, 101313. 10.1016/j.dcn.2023.101313

Nygaard, L. C., & Pisoni, D. B. (1998). Talker-specific learning in speech perception. Percept Psychophys, 60(3), 355–376. 10.3758/bf03206860

O’Sullivan, J. A., Power, A. J., Mesgarani, N., Rajaram, S., Foxe, J. J., Shinn-Cunningham, B. G., Slaney, M., Shamma, S. A., & Lalor, E. C. (2015). Attentional Selection in a Cocktail Party Environment Can Be Decoded from Single-Trial EEG. Cereb Cortex, 25(7), 1697–1706. 10.1093/cercor/bht355

Oostenveld, R., Fries, P., Maris, E., & Schoffelen, J. M. (2011). FieldTrip: Open Source Software for Advanced Analysis of MEG, EEG, and Invasive Electrophysiological Data. Computational Intelligence and Neuroscience, 2011. https://doi.org/Artn 156869 10.1155/2011/156869

Orf, M., Wostmann, M., Hannemann, R., & Obleser, J. (2023). Target enhancement but not distractor suppression in auditory neural tracking during continuous speech. iScience, 26(6), 106849. 10.1016/j.isci.2023.106849

Ortiz Barajas, M. C., Guevara, R., & Gervain, J. (2021). The origins and development of speech envelope tracking during the first months of life. Dev Cogn Neurosci, 48, 100915. 10.1016/j.dcn.2021.100915

Peltola, M. J., Leppanen, J. M., Maki, S., & Hietanen, J. K. (2009). Emergence of enhanced attention to fearful faces between 5 and 7 months of age. Soc Cogn ASect Neurosci, 4(2), 134–142. 10.1093/scan/nsn046

Quadrelli, E., Conte, S., Cassia, V. M., & Turati, C. (2019). Emotion in motion: Facial dynamics affect infants’ neural processing of emotions. Developmental Psychobiology, 61(6), 843–858. 10.1002/dev.21860

Reetzke, R., Gnanateja, G. N., & Chandrasekaran, B. (2021). Neural tracking of the speech envelope is differentially modulated by attention and language experience. Brain Lang, 213, 104891. 10.1016/j.bandl.2020.104891

Rigato, S., Stets, M., Charalambous, S., Dvergsdal, H., & Holmboe, K. (2023). Infant visual preference for the mother’s face and longitudinal associations with emotional reactivity in the first year of life. Sci Rep, 13(1), 10263. 10.1038/s41598-023-37448-8

Sai, F. Z. (2005). The role of the mother’s voice in developing mother’s face preference: Evidence for intermodal perception at birth. Infant and Child Development, 14(1), 29–50. 10.1002/icd.376

Souza, P., Gehani, N., Wright, R., & McCloy, D. (2013). The advantage of knowing the talker. J Am Acad Audiol, 24(8), 689–700. 10.3766/jaaa.24.8.6

Vonderlin, E., Ropeter, A., & Pauen, S. (2012). Erfassung des fruehkindlichen Temperaments mit dem Infant Behavior Questionnaire Revised. Zeitschrift fuer Kinder- und Jugendpsychiatrie und Psychotherapie, 40(5), 307–314.

Wagenmakers, E. J. (2007). A practical solution to the pervasive problems of p values. Psychon Bull Rev, 14(5), 779–804. 10.3758/bf03194105

Xie, W., McCormick, S. A., Westerlund, A., Bowman, L. C., & Nelson, C. A. (2019). Neural correlates of facial emotion processing in infancy. Dev Sci, 22(3), e12758. 10.1111/desc.12758

Yahav, P. H.-s., Sharaabi, A., & Golumbic, E. Z. (2024). The effect of voice familiarity on attention to speech in a cocktail party scenario. Cereb Cortex, 34, 1–16. 10.1093/cercor/bhad475

