## Supplementary material for "Neural Tracking of the Maternal Voice in the Infant Brain": SM2

*Table SM2. Central Face Encoding Accuracy. Exploratory Mixed Model analyzing potential differences in central face encoding. We found a main effect of VoiceFamiliarity and a main effect of Occipital_FaceEncodingAccuracy.*

### Mixed Model

Model Info

| Info | | |
|---|---|---|
| Model Type | Mixed Model | Linear Mixed model for continuous y |
| Model | lmer | Central_Faces_EncodingAccuracy_350_500_ms ~ 1 + VoiceFamilarity + Occipital_Faces_EncodingAccuracy_200_300_ms + Emotion + SessionOrder + Audio_EncodingAccuracy + VoiceFamilarity:Emotion + VoiceFamilarity:Occipital_Faces_EncodingAccuracy_200_300_ms + Emotion:Occipital_Faces_EncodingAccuracy_200_300_ms + VoiceFamilarity:Audio_EncodingAccuracy + Emotion:Audio_EncodingAccuracy + Occipital_Faces_EncodingAccuracy_200_300_ms:Audio_EncodingAccuracy + ( 1 \| SubjectID ) |
| Distribution | Gaussian | Normal distribution of residuals |
| Direction | y | Dependend variable scores |
| Optimizer | bobyqa | |
| DF method | Satterthwaite | |
| Sample size | 120 | |
| Converged | yes | |
| Y transform | none | |
| C.I. method | Wald | |

Note. Variable Occipital_Faces_EncodingAccuracy_200_300_ms is standardized

Note. Variable Audio_EncodingAccuracy is centered to the mean

#### Model Results

Model Fit

| Type | $R^2$ | df | LRT $X^2$ | p |
|---|---|---|---|---|
| Conditional | 0.481 | 12 | 66.804 | <.001 |
| Marginal | 0.405 | 11 | 65.736 | <.001 |

Additional Indices

| Info | Model Value | Comment |
|---|---|---|
| LogLikelihood | 303 | |
| AIC | -578 | Less is better |
| BIC | -539 | Less is better |

Fixed Effects Omnibus Tests

| | F | df | df (res) | p |
|---|---|---|---|---|
| **VoiceFamilarity** | 8.85991 | 1 | 90.9 | 0.004 |
| **Occipital_Faces_EncodingAccuracy_200_300_ms** | 44.50300 | 1 | 105.9 | <.001 |
| **Emotion** | 0.07552 | 1 | 80.1 | 0.784 |
| **SessionOrder** | 1.23986 | 1 | 81.8 | 0.269 |
| **Audio_EncodingAccuracy** | 2.78874 | 1 | 86.3 | 0.099 |
| **VoiceFamilarity ✻ Emotion** | 0.01812 | 1 | 80.2 | 0.893 |
| **VoiceFamilarity ✻ Occipital_Faces_EncodingAccuracy_200_300_ms** | 2.56028 | 1 | 106.6 | 0.113 |

Fixed Effects Omnibus Tests

| | F | df | df (res) | p |
|---|---|---|---|---|
| **Occipital_Faces_EncodingAccuracy_200_300_ms ✻ Emotion** | 0.86092 | 1 | 87.6 | 0.356 |
| **VoiceFamilarity ✻ Audio_EncodingAccuracy** | 0.03649 | 1 | 106.1 | 0.849 |
| **Emotion ✻ Audio_EncodingAccuracy** | 0.00247 | 1 | 80.8 | 0.961 |
| **Occipital_Faces_EncodingAccuracy_200_300_ms ✻ Audio_EncodingAccuracy** | 2.39931 | 1 | 108.0 | 0.124 |

Parameter Estimates (Fixed coefficients)

| | | | | 95% Confidence Intervals | | | | |
|---|---|---|---|---|---|---|---|---|
| **Names** | **Effect** | **Estimate** | **SE** | **Lower** | **Upper** | **df** | **t** | **p** |
| (Intercept) | (Intercept) | 0.03581 | 0.00165 | 0.03254 | 0.03909 | 30.9 | 21.6849 | <.001 |
| VoiceFamilarity1 | Stranger - Mother | 0.00802 | 0.00270 | 0.00268 | 0.01337 | 90.9 | 2.9766 | 0.004 |
| Occipital_Faces_EncodingAccuracy_200_300_ms | Occipital_Faces_EncodingAccuracy_200_300_ms | 0.00995 | 0.00149 | 0.00700 | 0.01291 | 105.9 | 6.6711 | <.001 |
| Emotion1 | happy - fearful | 6.81e-4 | 0.00248 | -0.00423 | 0.00559 | 80.1 | 0.2748 | 0.784 |
| SessionOrder1 | 2 - 1 | 0.00281 | 0.00252 | -0.00219 | 0.00781 | 81.8 | 1.1135 | 0.269 |
| Audio_EncodingAccuracy | Audio_EncodingAccuracy | 0.37621 | 0.22528 | -0.07043 | 0.82285 | 86.3 | 1.6700 | 0.099 |

Parameter Estimates (Fixed coefficients)

| Names | Effect | Estimate | SE | 95% Confidence Intervals Lower | 95% Confidence Intervals Upper | df | t | p |
|---|---|---|---|---|---|---|---|---|
| VoiceFamilarity1 ✻ Emotion1 | (Stranger - Mother) ✻ (happy - fearful) | -7.00e−4 | 0.00520 | -0.01102 | 0.00961 | 80.2 | -0.1346 | 0.893 |
| VoiceFamilarity1 ✻ Occipital_Faces_EncodingAccuracy_200_300_ms | (Stranger - Mother) ✻ Occipital_Faces_EncodingAccuracy_200_300_ms | 0.00448 | 0.00280 | -0.00107 | 0.01002 | 106.6 | 1.6001 | 0.113 |
| Occipital_Faces_EncodingAccuracy_200_300_ms ✻ Emotion1 | Occipital_Faces_EncodingAccuracy_200_300_ms ✻ (happy - fearful) | 0.00241 | 0.00259 | -0.00274 | 0.00755 | 87.6 | 0.9279 | 0.356 |
| VoiceFamilarity1 ✻ Audio_EncodingAccuracy | (Stranger - Mother) ✻ Audio_EncodingAccuracy | 0.08300 | 0.43454 | -0.77852 | 0.94452 | 106.1 | 0.1910 | 0.849 |
| Emotion1 ✻ Audio_EncodingAccuracy | (happy - fearful) ✻ Audio_EncodingAccuracy | 0.01779 | 0.35821 | -0.69240 | 0.72798 | 80.8 | 0.0497 | 0.961 |
| Occipital_Faces_EncodingAccuracy_200_300_ms ✻ Audio_EncodingAccuracy | Occipital_Faces_EncodingAccuracy_200_300_ms ✻ Audio_EncodingAccuracy | -0.41827 | 0.27003 | -0.95362 | 0.11709 | 108.0 | -1.5490 | 0.124 |

Random Components

| Groups | Name | Variance | SD | ICC |
|---|---|---|---|---|
| **SubjectID** | (Intercept) | 2.68e-5 | 0.00517 | 0.128 |
| **Residual** | | 1.83e-4 | 0.01353 | |

Note. Number of Obs: 120 , Number of groups: SubjectID 30

Random Effect LRT

| Test | N. par | AIC | LRT | df | p |
|---|---|---|---|---|---|
| **(1 \| SubjectID)** | 13.0 | -578 | 2.20 | 1.00 | 0.138 |

#### Estimated Marginal Means

Estimate Marginal Means - VoiceFamilarity

| | | | | 95% Confidence Intervals | |
|---|---|---|---|---|---|
| **VoiceFamilarity** | **Mean** | **SE** | **df** | **Lower** | **Upper** |
| Mother | 0.0318 | 0.00211 | 67.1 | 0.0276 | 0.0360 |
| Stranger | 0.0398 | 0.00216 | 67.9 | 0.0355 | 0.0441 |
