## Supplementary material for "Neural Tracking of the Maternal Voice in the Infant Brain": SM3

*Table SM3. Mixed Model analyzing ERP responses (Nc [400 – 800 ms, confirmatory], Nc [350 – 600 ms, exploratory], N290 [200 – 300 ms, confirmatory], and P400 [300 – 500 ms, confirmatory]. No significant effects of interest were found.*

Parameter Estimates (Fixed coefficients)

| | | | | 95% Confidence Intervals | | | | |
|---|---|---|---|---|---|---|---|---|
| **Names** | **Effect** | **Estimate** | **SE** | **Lower** | **Upper** | **df** | **t** | **p** |
| **Nc (400 – 800 ms)** | | | | | | | | |
| (Intercept) | (Intercept) | -4.6634 | 1.4993 | -7.641 | -1.6857 | 23.3 | -3.1104 | 0.005 |
| Trialnumber | Trialnumber | -0.1056 | 0.0662 | -0.237 | 0.0258 | 73.5 | -1.5964 | 0.115 |
| Emotion1 | fearful - ( happy, fearful ) | 0.3208 | 0.8320 | -1.332 | 1.9732 | 70.3 | 0.3856 | 0.701 |
| VoiceFamiliarity1 | Mother - ( Stranger, Mother ) | -1.1170 | 0.8754 | -2.856 | 0.6217 | 76.7 | -1.2759 | 0.206 |
| Session1 | 2 - ( 1, 2 ) | 3.9854 | 0.8762 | 2.245 | 5.7257 | 76.8 | 4.5484 | <.001 |
| Emotion1 ✻ VoiceFamiliarity1 | (fearful - ( happy, fearful )) ✻ (Mother - ( Stranger, Mother )) | -0.0803 | 0.8318 | -1.732 | 1.5718 | 70.3 | -0.0965 | 0.923 |
| **Nc (350 – 600 ms)** | | | | | | | | |
| (Intercept) | (Intercept) | -9.510 | 1.4470 | -12.384 | -6.6365 | 23.5 | -6.572 | <.001 |
| Trialnumber | Trialnumber | -0.109 | 0.0672 | -0.243 | 0.0245 | 66.2 | -1.622 | 0.110 |
| Emotion1 | fearful - happy | 0.657 | 1.7830 | -2.884 | 4.1980 | 70.6 | 0.368 | 0.714 |
| VoiceFamiliarity1 | Mother - Stranger | -3.060 | 1.8672 | -6.769 | 0.6481 | 77.0 | -1.639 | 0.105 |
| Session1 | 2 - 1 | 7.718 | 1.8687 | 4.007 | 11.4294 | 77.1 | 4.130 | <.001 |
| Emotion1 ✻ VoiceFamiliarity1 | (fearful - happy) ✻ (Mother - Stranger) | -0.629 | 3.5655 | -7.710 | 6.4528 | 70.6 | -0.176 | 0.861 |

Parameter Estimates (Fixed coefficients)

| | | | | 95% Confidence Intervals | | | | |
|---|---|---|---|---|---|---|---|---|
| **Names** | **Effect** | **Estimate** | **SE** | **Lower** | **Upper** | **df** | **t** | **p** |
| **N290 (200 – 300 ms)** | | | | | | | | |
| (Intercept) | (Intercept) | -9.510 | 1.4470 | -12.384 | -6.6365 | 23.5 | -6.572 | <.001 |
| Trialnumber | Trialnumber | -0.109 | 0.0672 | -0.243 | 0.0245 | 66.2 | -1.622 | 0.110 |
| Emotion1 | fearful - happy | 0.657 | 1.7830 | -2.884 | 4.1980 | 70.6 | 0.368 | 0.714 |
| VoiceFamiliarity1 | Mother - Stranger | -3.060 | 1.8672 | -6.769 | 0.6481 | 77.0 | -1.639 | 0.105 |
| Session1 | 2 - 1 | 7.718 | 1.8687 | 4.007 | 11.4294 | 77.1 | 4.130 | <.001 |
| Emotion1 ✻ VoiceFamiliarity1 | (fearful - happy) ✻ (Mother - Stranger) | -0.629 | 3.5655 | -7.710 | 6.4528 | 70.6 | -0.176 | 0.861 |
| **P400 (300 – 500 ms)** | | | | | | | | |
| (Intercept) | (Intercept) | 5.6407 | 1.9612 | 1.746 | 9.536 | 23.7 | 2.876 | 0.008 |
| Trialnumber | Trialnumber | -0.0451 | 0.0740 | -0.192 | 0.102 | 89.5 | -0.610 | 0.543 |
| Emotion1 | fearful - happy | -0.9276 | 1.6677 | -4.240 | 2.385 | 70.4 | -0.556 | 0.580 |
| VoiceFamiliarity1 | Mother - Stranger | -1.1780 | 1.7753 | -4.704 | 2.348 | 76.0 | -0.664 | 0.509 |
| Session1 | 2 - 1 | 6.2954 | 1.7772 | 2.766 | 9.825 | 76.1 | 3.542 | <.001 |
| Emotion1 ✻ VoiceFamiliarity1 | (fearful - happy) ✻ (Mother - Stranger) | 4.4459 | 3.3348 | -2.177 | 11.069 | 70.4 | 1.333 | 0.187 |
