## Supplementary material for "Neural Tracking of the Maternal Voice in the Infant Brain": SM4

*Table SM4. Correlation between difference in Nc response to happy and fearful faces and tracking accuracy. No significant effect of interest was found.*

Correlation Matrix

| | | audiotracking | Ncdifference |
|---|---|---|---|
| audiotracking | Pearson's r | — | |
| | df | — | |
| | p-value | — | |
| | 95% CI Upper | — | |
| | 95% CI Lower | — | |
| Ncdifference | Pearson's r | 0.239 | — |
| | df | 46 | — |
| | p-value | 0.102 | — |
| | 95% CI Upper | 0.490 | — |
| | 95% CI Lower | -0.049 | — |

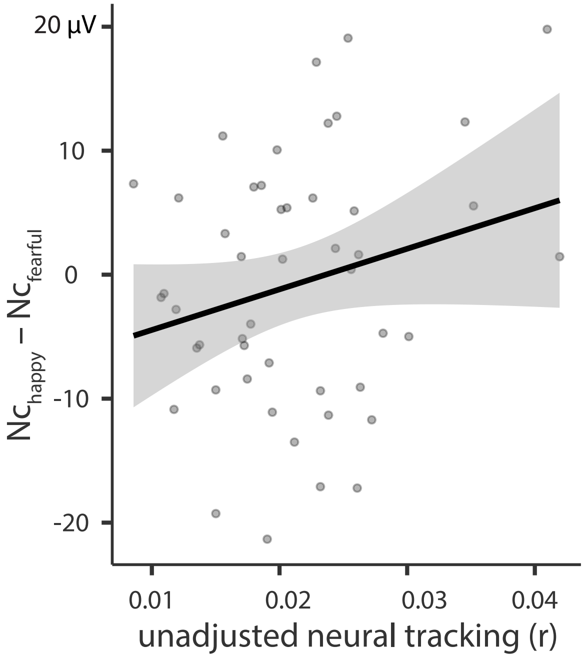
