## Supplementary material for "Neural Tracking of the Maternal Voice in the Infant Brain": SM1

*Table SM1. Central Auditory Encoding Accuracy. Confirmatory Mixed Model analyzing potential differences in central voice encoding. We found a main effect of VoiceFamiliarity.*

Parameter Estimates (Fixed coefficients)

| | | | | 95% Confidence Intervals | | | | |
|---|---|---|---|---|---|---|---|---|
| **Names** | **Effect** | **Estimate** | **SE** | **Lower** | **Upper** | **df** | **t** | **p** |
| (Intercept) | (Intercept) | 0.02170 | 0.00104 | 0.01963 | 0.02377 | 27.9 | 20.77956 | <.001 |
| VoiceFamiliarity1 | Stranger - Mother | -0.00445 | 9.41e-4 | -0.00631 | -0.00258 | 84.8 | -4.72225 | <.001 |
| Emotion1 | happy - fearful | 3.20e-6 | 9.26e-4 | -0.00183 | 0.00184 | 84.4 | 0.00345 | 0.997 |
| Central_Faces_EncodingAccuracy | Central_Faces_EncodingAccuracy | -0.02135 | 0.03038 | -0.08155 | 0.03885 | 98.6 | -0.70264 | 0.484 |
| Log BaselinePower | Log BaselinePower | 3.49e-4 | 1.12e-4 | 1.27e-4 | 5.71e-4 | 106.3 | 3.11329 | 0.002 |
| VoiceFamiliarity1 ✻ Emotion1 | (Stranger - Mother) ✻ (happy - fearful) | -7.20e−5 | 0.00186 | -0.00375 | 0.00360 | 84.4 | -0.03878 | 0.969 |
